# How bats exit a crowded colony when relying on echolocation only - a modeling approach

**DOI:** 10.1101/2024.12.16.628648

**Authors:** Omer Mazar, Yossi Yovel

## Abstract

Bats face a complex navigation challenge when emerging from densely populated roosts, where vast numbers take off at once in dark, confined spaces. Each bat must avoid collisions with walls and conspecifics while locating the exit, all amidst overlapping acoustic signals. This crowded environment creates the risk of acoustic jamming, in which the calls of neighboring bats interfere with echo detection, potentially obscuring vital information. Despite these challenges, bats navigate these conditions with remarkable success. Although bats have access to multiple sensory cues, here we focused on whether echolocation alone could provide sufficient information for orientation under such high-interference conditions. To explore whether and how they manage this challenge, we developed a sensorimotor model that mimics the bats’ echolocation behavior under high-density conditions. Our model suggests that the problem of acoustic jamming may be less severe than previously assumed. Frequent calls with short inter-pulse intervals (IPI) increase the sensory input flow, allowing integration of echoic information across multiple calls. When combined with simple movement-guidance strategies—such as following walls and avoiding nearby obstacles—this accumulated information enables effective navigation in dense acoustic environments. Together, these findings demonstrate a plausible mechanism by which bats may overcome acoustic interference and underscore the role of signal redundancy in supporting robust echolocation-based navigation. Beyond advancing our understanding of bat behavior, they also offer valuable insights for swarm robotics and collective movement in complex environments.

## Introduction

In many bat species individuals dwell together in caves (or similar roosts), forming large colonies with tens to several millions of individuals^1,2^. Each evening, at approximately the same time, the bats take off from their roost, navigating through its passages toward the exit. The high density of bats flying simultaneously in great proximity poses many challenges for orientation in such a crowded and noisy environment. Flying while avoiding collisions, often in a pitch-black cave, demands the continuous detection and localization of both obstacles and nearby bats^3,4^. Employing echolocation, bats emit strong ultrasonic signals and interpret the reflected echoes to perceive their surroundings^5^. The reception of neighbors’ loud calls, which share similar acoustic features with their own calls, can potentially hinder the bats’ ability to detect the faint echoes reflected off the walls and the surrounding bats^5,6^. We examined whether bats could rely solely on echolocation to exit the roost even during such a chaotic ‘rush hour’.

The question of how bats cope with acoustic interference — i.e., the masking of potential echoes by conspecific signals — has been extensively researched using playback experiments, field observations, on-body tags, and computational simulations^7–17^. However, much of this research has focused on foraging bats in small groups^5,6,9,16,18–20^. The challenges bats encounter during roost exits (e.g., cave exits) differ markedly from those encountered during group foraging. Bat density during roost exits is significantly higher, and bats need to detect and follow static walls or obstacles, which produce loud echoes, rather than small, sporadic prey items that generate faint echoes^21^. Their flight during exits is also more directional and involves avoiding collisions with conspecifics, in contrast to the erratic hunting maneuvers typically observed while foraging. Echolocation studies during dense collective movement are scarce^4,6,22–25^, likely due to the complexities in recording separate echolocation calls and tracking individual flights within the swarm.

While collective movement has been extensively studied in various species, such as insect swarming, fish schooling, and bird murmuration^26–32^, as well as in swarm robotics, where agents perform tasks such as coordinated navigation and maze-solving^33–35^, most studies have focused on movement algorithms that assume full detection of neighbors^36–43^. Some models have incorporated limited interaction rules where individuals respond to only one or a few neighbors due to sensory constraints^44,45^ However, fewer studies have explicitly examined how sensory interference, occlusion, and noise influence decision-making and affect collective movement^46^.

The present study addresses these gaps by introducing an agent-based sensorimotor model based on the well-documented echolocation capabilities of bats, simulating multiple bats pathfinding their way out of a cave-like structure. We modeled the echolocation behavior of two insectivorous bat species: *Pipistrellus kuhlii (*PK), which roosts in abandoned buildings and frequently navigates through conspecific-dense, cluttered corridors and the cave dwelling *Rhinopoma microphyllum* (RM) which emerges from its roosts with thousands of individuals simultaneously. These two species differ in their echolocation signals - PK echolocation signals are characterized by a wider bandwidth and a higher terminal frequency than RM calls. We quantified the performance of an individual bat flying among conspecifics, demonstrating that even a relatively simple sensorimotor algorithm can facilitate successful orientation in such complex environments. The modeling approach enabled us to explore how various biological and ecological factors influence successful navigation under such challenging conditions.

## Results

Our model was designed with conservative assumptions regarding bats’ sensing, movement, and sensorimotor integration, aiming to underestimate their capabilities and thereby establish a lower bound on their actual performance. Real bats likely outperform the model’s predictions. In our 2D simulations^7^, each bat emits sound signals and receives echoes reflected from the roost walls and other bats, while also encountering masking signals caused by calls from conspecifics. These masking signals can interfere and completely eliminate echo detection (which we refer to as jamming) or cause echo localization errors. After estimating the distance and direction of each detected reflector, the bat adjusts its echolocation parameters and maneuvers to find the exit while simultaneously avoiding collisions. The bats dynamically adjust their echolocation parameters—including call rate, duration, and frequency range—based on the estimated distance to obstacles, following the well-documented transition between search, approach, and buzz phases observed in echolocating bats (see^7^ and Methods). Their reception was modeled using a biologically inspired filter-bank receiver comprising 80 gammatone channels^7,47,48^. Each bat adjusted its flight following a simple pathfinding algorithm based solely on the estimated locations of the detected reflectors (see Methods, Supplementary Figure 1, and Supplementary Movie 1 for additional details). The bats had to exit a roost designed as a corridor (14.5 m long x 2.5 m wide), with a right-angle turn located 5.5 m before the exit (Figure 1A). Additionally, an obstacle (1.25 m wide) was situated 2.25 m in front of the opening. The simulated bats initiated their flight from the far end of the corridor, within a randomly selected 1.5 × 2 m² area, taking off in the general direction of the exit (±30 degrees), without prior knowledge of the roost’s structure.

**Figure 1:**
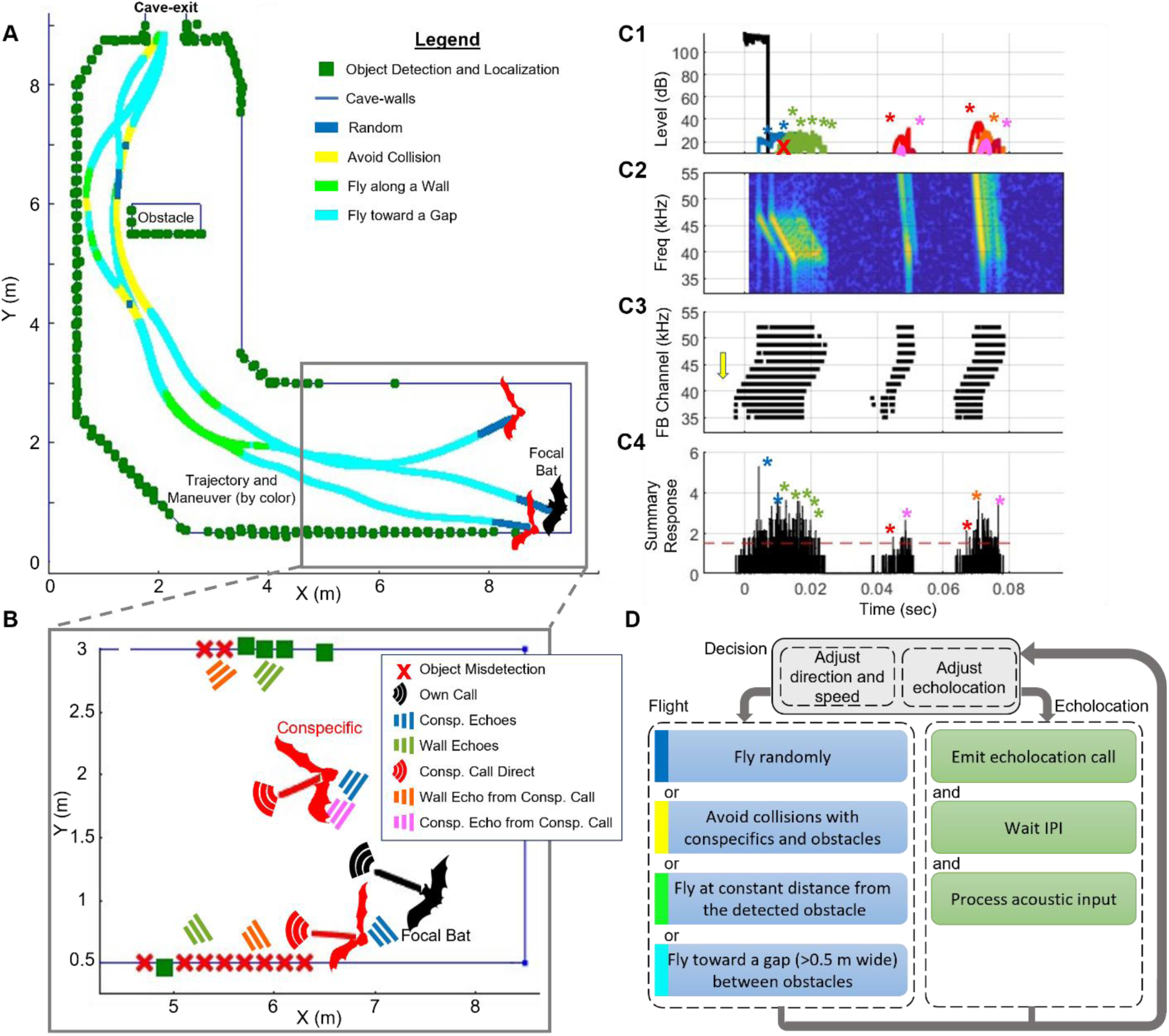
The sensorimotor model. **(A)** Top view of the cave with three bats’ trajectories. The focal bat is shown in black. All bats’ flight trajectories are displayed while the bats’ moment-to-moment decisions are represented by the colored lines: blue - random flight, yellow – collision avoidance, light green – wall-following, turquoise – movement toward a wall gap (see panel D for details). Green squares depict reflectors detected by the focal bat along its route. **(B)** A zoomed-in view of the marked rectangular area in Panel A, where the focal bat (black) emitted one echolocation call (black) and received echoes from the cave walls (green) and from two other bats (blue). It also received conspecifics’ calls (red) and their reflection from the cave walls (orange), as well as the reflections from other bats (pink). Green squares indicate points that were detected by the focal bat from this call and red x’s indicate missed points due to acoustic masking (i.e., jammed reflectors). The locations of the detected reflectors (green squares) are marked according to their localization by the bat (with simulated errors). The lines near the bats depict their flight direction. **(C)** The acoustic scene received by the focal bat is as depicted in B, including the emitted call and all received signals (colors as in panel B). **(C1)** The time-domain plot displays the envelope of signals, encompassing the emitted call and the received signals: the desired echoes from the walls and conspecifics; the calls of other bats; the echoes returning from conspecific calls and reflected off the walls and off other bats. Notably, in this example, some of the desired wall-echoes are jammed by stronger self-echoes reflected from nearby conspecifics. **(C2)** The spectrogram of all the received signals presented in C1; for clarity, the emitted call is not depicted. **(C3)** The responses of the active channels of the cochlear filter bank (FB channel) after de-chirping. Each channel is represented by its central frequency on the y-axis. Each black dot represents the timing of a reaction that was above the detection threshold in each channel. Note that early reactions in low-frequency channels (marked by a yellow arrow) result from the stimulation of those channels caused by the higher frequencies of the downward FM chirp. However, most of these stimulations do not reach the detection threshold and are therefore not detected (see Methods). **(C4)** The detections of each channel are convolved with a Gaussian kernel, summed, and compared with the detection threshold (dotted red line). Colored asterisks mark peaks that were classified as **successful detections**—those identified in both the interference-free and full detection conditions (see Methods for details). Other peaks may originate from masking signals or overlapping echoes that did not meet the detection criteria (colors of the sources are as defined above). **Panel D** depicts the pathfinding algorithm used by the bat. The algorithm involves a correlated-random flight during the search phase (blue), collision avoidance (yellow), flying along the wall at a constant distance (green), and flying toward the center of a gap between obstacles as an indicator of a possible exit (cyan). After each echolocation call, the bat awaits an IPI (Inter Pulse Interval) period before processing the detections, adjusting flight and echolocation parameters, and emitting the next call. Based on the received signals, it then modifies its next call design and adjusts its direction and speed accordingly. For a detailed diagram of the complete sensorimotor process see Supplementary Figure 1.

The sensory model accounted for six types of acoustic signals: (1) the bat’s own calls, (2) echoes from conspecifics, (3) echoes from walls in response to the bat’s own calls (i.e., desired wall echoes), (4) echoes from conspecific calls reflected off other bats, (5) echoes from conspecific calls reflected off walls, and (6) the conspecific calls themselves. In the baseline model, bats were assumed to reliably distinguish between all these signal types. In contrast, the confusion model described below specifically tested the impact of failing to distinguish between desired wall echoes (3) and wall echoes generated by conspecific calls (5), while preserving the bat’s ability to identify all other signal types. In brief, the bat responded to echoes as follows (see Methods and Supplementary Figure 1 for details): If an obstacle or a conspecific was detected in front of the bat and was too close, the bat would maneuver to avoid a collision. Otherwise, for exit-seeking, the bat would follow the contour of the walls by steering toward the farthest detected obstacle ahead. If a gap greater than 0.5 m was identified between adjacent reflectors, the bat directed its trajectory toward the center of the gap.

The ability of the bats to exit the roost within 15 sec was evaluated for different group sizes, from a single bat and up to 100 individuals. For simplicity, we will refer to the initial density at the cave’s far end as the number of bats per 3m^2^ (i.e., for groups of 100 bats, the density is 100 bats/3m^2^, or 33.3 bats/m^2^). The bat densities we tested were chosen to reflect the typical range of bat densities observed in natural caves during emergence events^25,49,50^. Key model parameters, such as the sensory integration window, object target strength, echolocation parameters, and flight velocity (see Table 1), were manipulated and their impact on the exit performance was analyzed. To explicitly quantify the effect of sensory masking vs. the effect of collision avoidance (i.e., spatial interference) only, we turned the acoustic interference on and off to measure its impact. Each scenario was repeated as follows: 1 bat: 240; 2 bats: 120; 5 bats: 48; 10 bats: 24; 20 bats: 12; 40 bats: 12; 100 bats: 6 (see Table 1), while misidentification rate, multi-call clustering, and wall/conspecific target strength were tested only up to 40 bats (see Table 1).

**Table 1:** Key model parameters and their effects on performance metrics. The table presents the key parameters tested, their ranges, default values, and effect sizes on various performance metrics: exit probability, time-to-exit, jamming probability, and collision rate with obstacles. The parameters comprised the number of bats, bat species (PK-*Pipistrellus kuhlii*, RM –*Rhinopoma microphyllum*), integration window, nominal flight speed, call level, echo mis-identification with multi-call clustering (yes/no), masking (yes/no), wall target strength, and conspecific target strength. In each scenario, all parameters except the tested one were set to the default value. Call levels are reported in dB-SPL, referenced at 0.1 m from the source. Effect sizes for each parameter are explicitly listed for all four-performance metrics, expressed as the change per unit of the tested parameter (e.g., per bat or per 10 dB). For flight speed, a non-monotonic relationship was observed, and values are reported both before and after the peak performance (see Results, Fig. 3B).Values in square brackets indicate the minimum and maximum of the metric across the tested range.. Asterisk (*) indicates a significant impact. Each scenario was tested using Generalized Linear Models (GLMs) with number-of-bats and the tested parameters set as fixed explaining variables. Exit probability and jamming probability were treated as binomially distributed, collision rate was treated as a Poisson distributed, and all other variables were considered normally distributed. Explaining variables were set as fixed factors. The number of repetitions for each scenario was as follows: 1 bat: 240; 2 bats: 120, 5 bats: 48; 10 bats: 24; 20 bats: 12; 40 bats: 12; 100 bats: 6. ^Ω^ Misidentification rate, multi-call clustering, wall target strength, and conspecific target strength were simulated only up to 40 bats due to significantly longer run-times. ^ѱ^ A significant difference in call intensity was found only for a bat density of 100 bats/3m^2^, and between the group with a level of 100dB-SPL and all other groups. ^α^ see Supplementary Figure 3. ^β^ see Supplementary Figure 4.

| Key parameter | Tested range | Default value | Effect Size on Explained Variable |  |  |  | GLM Explaining factors |
| --- | --- | --- | --- | --- | --- | --- | --- |
|  |  |  | Exit prob. (%) | Time-to-exit (sec) | Jamming-prob. (%) | Obs. Collision (sec<sup>-1</sup>) |  |
| Number Of Bats<sup>Ω</sup> | [1, 2, 5, 10, 20, 40, 100] | **All Values** | -0.37/bat*<br>[63:100] | 0.044/bat *<br>[4.6:8.9] | 0.54/bat*<br>[0:54] | 0.25/bat*<br>[0.05:0.3] | Number of bats |
| Bat species | [PK, RM] | **PK** | 4.5<br>[63:67] | -0.4*<br>[8.5:8.9] | 9*<br>[54:63] | 0.01<br>[0.29:03 ] | Number of bats, bat species |
| Integration Window (#) | [0,1,3,5,10] | **5** | 0.41/call*<br>[29:70] | -0.16/call*<br>[9.2:7.6] | 0.01/call<br>[54:55] | 0.27/call<br>[0.25:0.53] | Number of bats, integration window size |
| Nominal flight speed (m/s) | [2,4,6,8,10] | **6** | 6/(1m/s),<br>-12/(1m/s)*<br>[15:63] | -1/(1m/s),<br>1/(1m/s)*<br>[6:9.6] | 1.4/(1m/s)*<br>[55:65] | 0.18/(1m/s)*<br>[0.07:1.3] | Number of bats, flight speed, square of flight speed |
| Call level (dB-SPL, @ 0.1m) | [100,110,120, 130] | **120** | 13<sup>ψ</sup><br>[50:63] | -0.5<br>[7.9:8.4] | 0.5/10dB*<br>[53:58] | 0.07 <sup>ψ</sup><br>[0.29:0.36] | Number of bats, call level |
| Misidentification <sup>Ω</sup> | [Yes/No] | **N** | -69*<br>[14:83] | 1.3*<br>[7.6:8.9] | 30*<br>[50:80] | 0.6*<br>[0.2:0.8] | Number of bats, With and without confusion |
| Misidentification and multi-call clustering <sup>Ω</sup> | [Yes/No] | **N** | -23*<br>[58:83] | 1.7*<br>[7.6:9.3] | 29*<br>[50:79] | -0.02*<br>[0.18:0.2] | Number of bats, With and without multi-call clustering |
| Masking | [Yes/No] | **Y** | 23*<br>[63:86] | 0.8*<br>[8.0:8.8] | 54*<br>[0:54] | 0.03<br>[0.26:0.29] | Number of bats, With and without masking |
| Wall target strength (dB) <sup>α, Ω</sup> | [-33, -23, -13, -3] | **-23** | 16/10dB *<br>[23:87] | 0.7/10dB *<br>[6.6:9.5] | -8.5/10dB *<br>[34:68] | 0.07/10dB *<br>[0.19:0.47] | Number of bats, wall target strength |
| Conspecific target strength (dB) <sup>β, Ω</sup> | [-49, -43, -33, -23] | **-23** | -1.5/10dB<br>[85:91] | 0.25/10dB*<br>[7.2:8.15] | -0.5/10dB<br>[48:50] | 0.1/10dB<br>[0.16:0.2] | Number of bats, conspecific target strength |

### Bats find their way out of the cave even at high conspecific densities

We first examined how bat density affects bats’ ability to exit the cave, both alone and in a group. The probability of exiting the cave within 15 seconds—defined as the proportion of bats that successfully exited within this time frame—was significantly reduced at higher densities (Figure 2A, see Supplementary Movie 1 for a view of the bats’ movement, p<10^-10^, t =-23, DF=4077, GLM, see details in Table 1). In trials in which a single bat was flying alone, it successfully exited the cave in 100% of the cases. Even without sensory interference, the probability of exiting decreased significantly from 100% to 86%±1.4% and 91%±1.7% at densities of 100 PKs/3m^2^ and 100 RMs/3m^2^, respectively (mean ± s.e.). When acoustic interference was added, the exit probability further decreased to 63%±1.4% and 67%±1.4% for 100 PKs and RMs, respectively (see Figure 2A).

**Figure 2:**
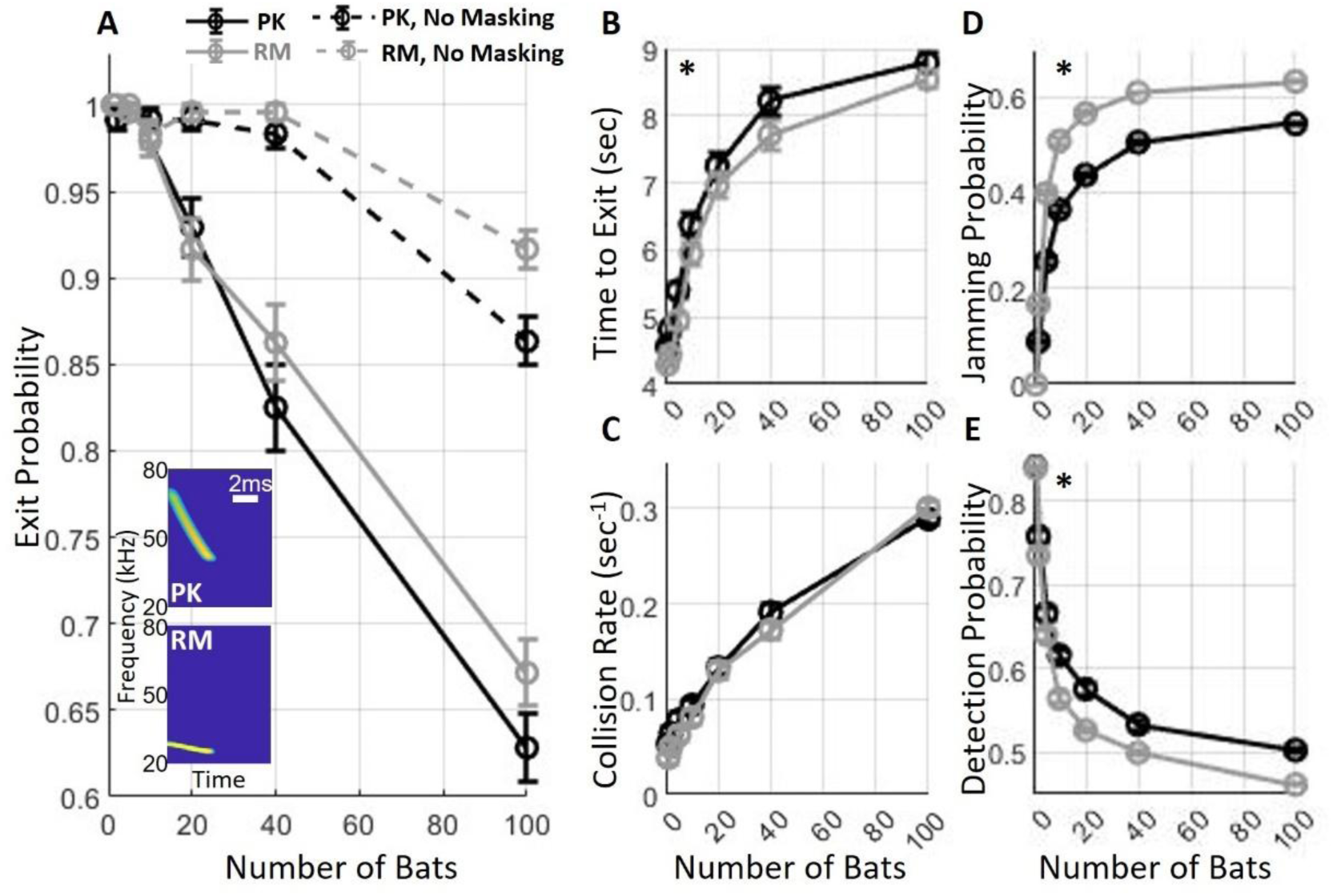
Exit performance of *P. Kuhlii* (PK) and *R. Microphyllus* )RM(. **(A)** Sensory interference significantly impaired the probability of exiting the cave (compare dashed lines with continuous lines). The probability of a successful exit also declined as the number of bats increased, with no significant difference observed between the species when masking interference was applied. The insert shows the spectrograms of the echolocation calls of PK (top) and RM (bottom). **(B)** The time-to-exit, which was calculated for successful trials only, and **(C)** the collision rate with the walls both increased as a function of the number of bats. **(D)** The probability of jamming significantly increased to about 55% and 63% with 100 bats for PK and RM, respectively. **(E)** The detection probability of a wall reflector at one meter or less in front of a bat decreased as a function of the number of bats. In panels (A-E), circles represent means and bars represent standard errors (see details in Table 1). Asterisks indicate significant differences between the lines in each panel.

The difference in exit probability between the two species was not significant (p=0.08, t =1.74, DF=4077, GLM as above, Figure 2A). Similarly, the difference in echolocation parameters between the two species did not affect the collision rate with the walls (with a maximum of 0.29 and 0.3 collisions per bat per second for PK and RM, respectively, with 100 bats (p=0.63, t =-0.48, DF=4077, GLM, Figure 2C, see details in Table 1). To quantify sensory interference, we defined a jammed echo as an echo entirely missed due to masking. The jamming probability, which was calculated as the number of jammed echoes divided by the total number of self-echoes, was significantly higher for RM compared to PK. The maximum difference between the two models was 14.3% at a density of 10 bats, with a smaller difference of 9.8% observed at 100 bats (p<10^-10^, t =6.56, DF=4077, GLM, Figure 2D, see details in Table 1). Accordingly, PK demonstrated a minor but significant advantage in detecting the cave walls (p=0.024, t =-2.25, DF=4077, GLM, Figure 2E, see details in Table 1). With 100 bats flying together, the probability of detecting a wall echo at a distance of 1 m in a single call was around 50% and 46% for PK and RM, respectively. Despite this minor disadvantage in detection, RM bats exhibited a better time-to-exit average than PK bats, being 0.5 seconds faster to exit (p=0.0005, t =-4.06, DF=3533, for n=40 bats, Figure 2B). Additionally, RM bats experienced a significantly higher probability of their self-generated echoes, reflected off conspecifics, being jammed (p = 0.00016, t = 3.8, DF = 3593, GLM; see details in Table 1).

### Multi-call integration improves exit performance

We next examined whether bats improve their performance when integrating information from several consecutive calls. The integration window determines the number of previous calls the bat uses at each step to guide its next movement decision (see Methods and Supplementary Figure 2A). In the basic multi-call integration model, detections from the previous calls — by default the last five — were stored in an allocentric (x-y) reference frame, with each detection treated independently as a potential obstacle without clustering or filtering. At each decision, the bat takes all of these detections into account when guiding its movement and echolocation. The probability of exiting the roost significantly increased when increasing the size of the integration window for all bat densities (p<10^-10^, t =28.5, DF=10197, GLM, Figure 3A, see details in Table 1). For example, at a density of 40 bats/3m^2^, the exit probability improved from 20%, to 75%, and to 87% as the window size increased from one, to three, and to 10 previous calls, respectively. In addition, increasing the window size resulted in a significant improvement in the time-to-exit and the avoidance of wall collisions (p<10^-10^, t =-12.8, DF=7661; p<10^-10^, t =-46.5, DF=10197, respectively, GLM, see details in Table 1). With 100 bats, the collision rate decreased by a factor of 2 from 0.53 to 0.25 collisions per second as the window increased from 1 to 10 calls. The size of the integration window had no significant effect on the jamming probability (p=0.37, t =0.9, DF=10197, GLM, see details in Table 1).

**Figure 3:**
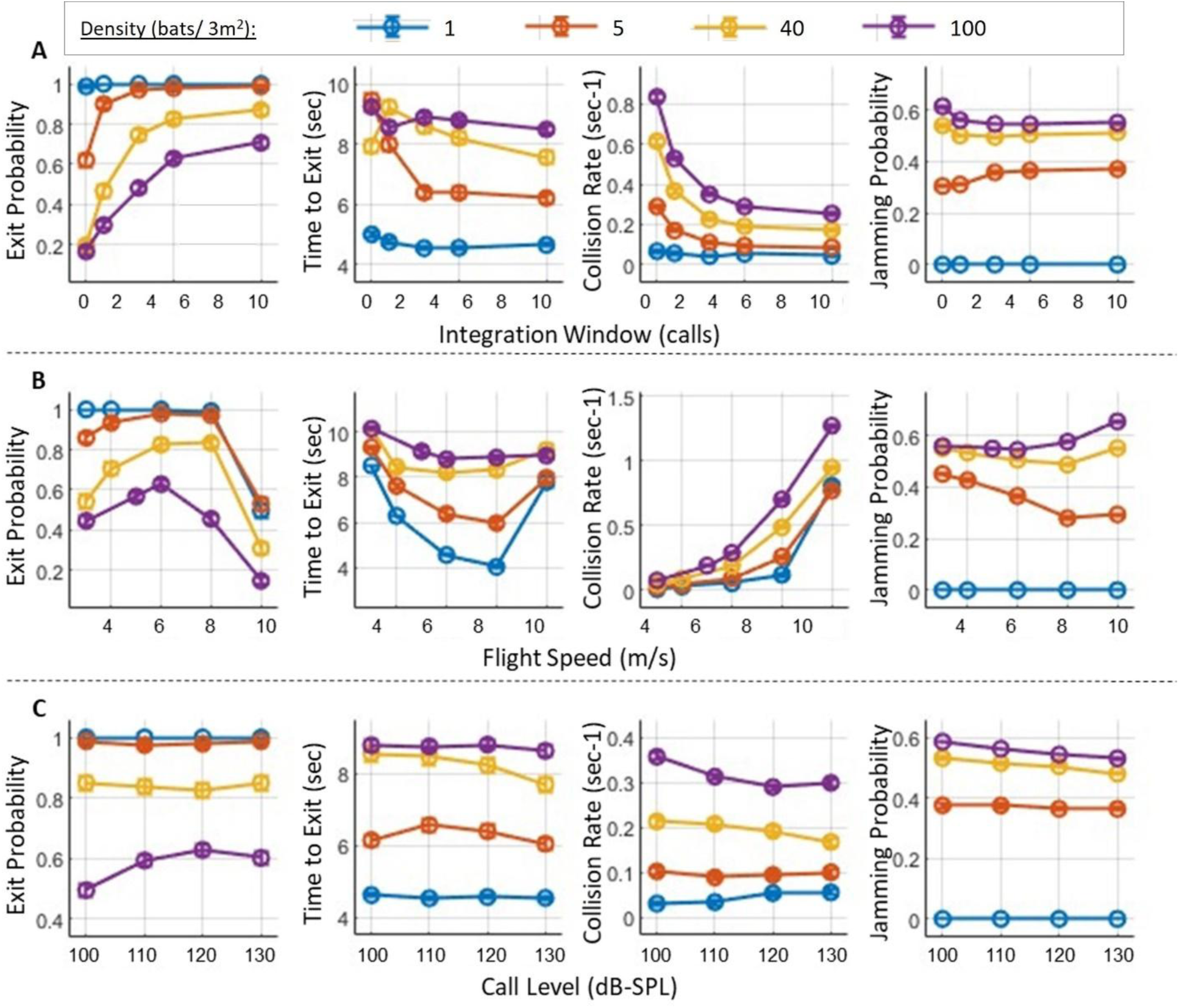
Exit performance as a function of key sensorimotor parameters. **(A)** The effect of the integration window on the probability of exiting the roost, the time-to exit, the rate of collisions with the walls, and the probability of jamming (from left to right, respectively). Each colored line shows the trend as a function of the window-size for different bat densities, with each color representing a specific density. Note that a window size of 0 indicates that only the most recent call is used in the bat’s decision-making, without integrating detections from previous calls**. (B)** The effect of the nominal flight speed of the bats, with panels and line-colors as in panel A. An optimal speed of approximately 6 to 8 m/sec can be observed for all densities above one bat. **(C)** The effect of call intensity on exit performance, panels as in (A). In all panels, circles represent means and bars represent standard errors. Error bars depicting standard errors are presented but are very small due to the large number of simulation repetitions. See Table 1 for the number of simulated bats.

### Exit probability was maximal at an intermediate flight-speed

We observed a significant and non-linear effect of the flight speed of the bats on the performance, as shown in **Figure 1 Figure 3B** (p<10^-10^, t =-29.9, DF=10196, GLM, see details in Table 1). The exit probability increased with flight speed until it reached a maximum at 6-8 m/s and then declined rapidly. This was the case for all bat densities, with the maximal exit probability ranging between 65% to 99%. At the optimal velocity, the time-to-exit was also minimal. However, the collision rate increased monotonically with speed, with a steep incline above the optimal speed.

### Call intensity had only a minor effect on exit performance and only at high bat densities

For low bat densities (<40 bats), call intensity did not have a significant impact on either exit probability or collision rate (Figure 3C, p=0.89, t =0.13, DF=5757; p=82, t -0.21, DF=5757, respectively, GLM, see details in Table 1). Call intensity affected exit performance only when the intensity dropped to 100 dB-SPL (@ 0.1m) and only at a high bat density of 100 bats/3m^2^ (Figure 3**Figure 3** C). In this scenario, the exit probability declined from approximately 60% to 49.5% (p=0.003, F = 8.45, DF = 2396, One-way ANOVA with ’hsd’ post hoc test), and the collision rate increased from 0.3 to 0.35 collisions per second (p<3·10^-6^, F = 22.18, DF = 2396). Notably, this low intensity is below the typical search-call intensity of most echolocating bats. At the same bat density (100 bats/3m^2^), further increasing the call intensity to above 100dB-SPL had no significant effect on either exit probability (p=0.6) or collision rate (p=0.07). Calling louder also slightly, but significantly, decreased the jamming probability at all bat densities, with a decrease of 3.5%±8% to 5.5%±5% (mean ± s.e.) (p=0.02, t =-2.26, DF=8157, GLM, see Table 1).

### While confusion between the desired echoes and those from conspecific calls may significantly impair exit performance, multi-call clustering helps to mitigate this

We next addressed the challenge of echo classification, assuming that a bat can differentiate an echo resulting from its own calls from echoes resulting from the calls of other bats. To examine this assumption, we tested another model, referred to as the **confusion model**, in which bats responded similarly both to wall echoes returning from their own emissions and to those from conspecific emissions, treating all as their own echoes. This confusion significantly decreased exit performance for all bat densities (above one bat). The probability of a successful exit for a density of 40 bats/3m^2^ dropped from 83.3±2.4% to 14.6±2.3% (p<<10^-10^, t =-20.7, DF=2877, GLM, see details in Table 1), the exit time increased from 7.6±0.18 to 9.3±0.2 seconds (p<<10^-10^, t =15.5, DF=2157, GLM), and the collision rate increased significantly from 0.2±0.007 to 0.8±0.013 collisions per second (p<<10^-10^, t =-30, DF=28777, GLM, see Figure 4, red and yellow lines).

**Figure 4:**
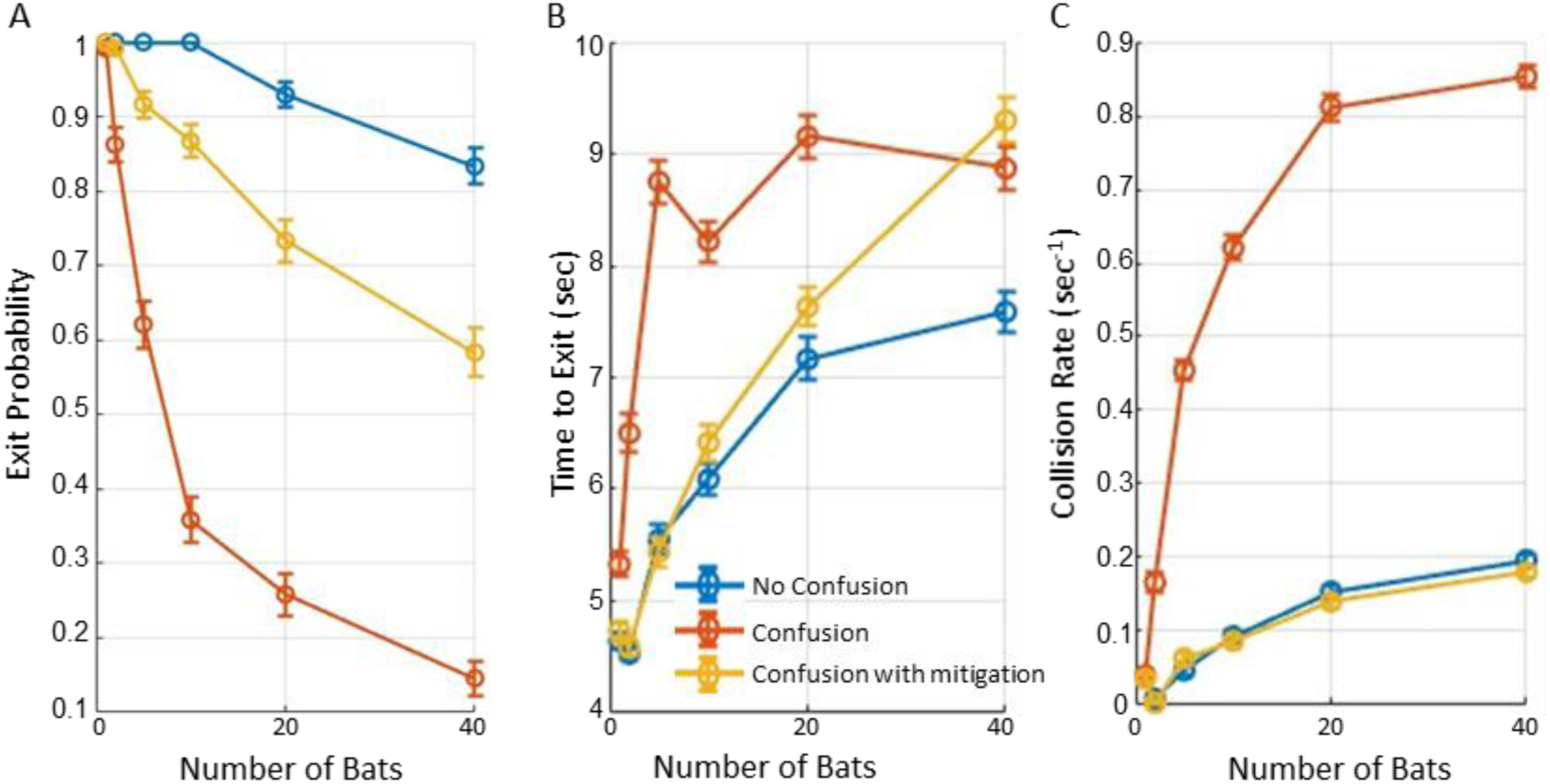
The impact of confusion on performance. The figure illustrates the impact of classification confusion on roost-exit performance under various conditions. Blue lines depict trials with masking, while assuming that bats can distinguish between echoes from their own calls and those of conspecifics (referred to as "No Confusion"). Red lines depict performance where confusion between echoes is assumed. Yellow lines depict performance under the confusion condition, with the added capability of multi-call clustering in a short-term working memory (referred to as “confusion with mitigation”, see text for further details). In all panels, circles represent means and bars indicate standard errors. **(A)** The probability of exiting the roost significantly decreased with masking and confusion. In conditions with confusion and no aggregation process, only 15% of bats successfully exited the roost, at a density of 40 bats/3m^2^. Multi-call clustering partially mitigated the confusion effect but did not eliminate it. **(B)** Bats with the ability to distinguish between echoes demonstrated significantly shorter exit times than those experiencing confusion. Note that time-to-exit refers only to successful attempts. **(C)** The collision rate with walls was highest for bats experiencing both masking and confusion but decreased significantly when without confusion. Multi-call clustering restored performance to the "No Confusion" condition, reducing collision rates accordingly, at densities between 1 to 40 bats/3m^2^.

To further examine whether this substantial decrease in performance could be mitigated even without improving echo identification, we tested an enhanced integration model that, in addition to extending the number of calls integrated, clustered spatially close detections, removed outliers, and estimated wall directions based on grouped reflectors (see Methods and Supplementary Figure 2B). This **’multi-call clustering’** significantly improved performance, but exit probability and time-to-exit still remained significantly lower than without echo-confusion: exit probability = 58±3% in comparison to 83.3±2.4% without echo confusion (p<<10^-10^, t =18.3, DF=28777, GLM), time-to-exit =9.3±0.2 seconds (p<<10^-10^, t =-13.7, DF=1996, GLM), see Figure 4, yellow line. The results above are reported for a density of 40 bats/3m^2^. Interestingly, the multi-call clustering restored the collision rate to the levels observed under the "No Confusion" condition (p=0.68, t =-0.42, DF= 2877, GLM, see Figure 4C, dark-purple and red lines).

#### Effect of Wall and Conspecific Target Strengths on Exit Performance

Increasing the wall target strength significantly enhanced navigation performance (Supplementary Figure 3, Table 1), improving exit probability by up to 64% and reducing time-to-exit by up to 2.8 seconds (p << 10⁻¹⁰). Stronger wall echoes improved environmental awareness but also slightly increased masking of desired conspecific signals.

In contrast, changes in conspecific target strength had a much smaller effect (Supplementary Figure 4, Table 1), with only minor improvements in detection and collision rates, and no significant impact on exit probability. This likely reflects the fact that both desired and masking signals scale similarly with conspecific reflectivity. Overall, the model showed low sensitivity to variations in conspecific target strength.

## Discussion

We present a model-based approach that suggests how echolocating bats might find their way out of a crowded roost while contending with severe sensory interference caused by numerous nearby conspecifics. Our results demonstrate that a single bat, lacking prior knowledge of the roost’s structure, successfully found the exit in all simulated trials using echolocation alone. As bat density increases, the bats face increased collision risks and more substantial acoustic interference, both of which reduce the probability of efficiently finding the exit. Nevertheless, even at densities of 100 bats/3m^2^, most bats (63%) successfully exited the roost within a short timeframe. These results are based on a 2D simulation with up to 33 bats/m², under the assumption that bats can distinguish their own echoes from those of conspecifics. We demonstrate how a simple sensorimotor approach can solve this supposedly challenging task. This approach encompasses the following principles: (1) emission of echolocation calls; (2) reception of reflected echoes and masking signals; (3) detection of reflectors (including walls and conspecifics) using a gammatone filter bank biological receiver; (4) localization of the detected objects; (5) employment of multi-call integration of acoustic detections; (6) adjustment of flight and echolocation behavior based on the distance and angle to the reflectors; and (7) application of simple pathfinding rules to follow walls and gaps while avoiding collisions. Notably, despite the jamming of a substantial percentage of the echoes — particularly, with 100 bats, 50% of the echoes from nearby obstacles at ∼1 m distance — the bats managed to maneuver correctly even with this simple approach and partial data.

A key component of this success was the multi-call integration: increasing the number of stored calls from one to ten markedly improved performance, raising the exit probability from 20% to 87% and halving the collision rate. Real bats likely use a much more sophisticated approach that also includes memorizing the roost’s structure^51^, using landmarks inside the roost^52^, reliance on the movement of nearby conspecifics^43,49^, and exploitation of other sensory modalities. We thus expect their actual performance to surpass that of our modeled bats.

Our model suggests that acoustic jamming might be less problematic than has been generally assumed^5,11,53^, and that movement under severe acoustic masking could be mitigated by increasing the call-rate, creating a redundancy across several calls-similar to how real bats behave in a complex environment^6^. In our model, the Inter-Pulse Interval (IPI) naturally varied according to established echolocation behavior, decreasing from 100 msec in the search phase to 35 msec (∼28 calls per second) in the approach phase, and further to 5 msec (200 calls per second) during the final buzz (Table 2). The results indicate that this redundancy, combined with simple sensorimotor heuristics, enhances successful navigation. This is consistent with several recent studies that have pointed in this direction^7,24,25^.

**Table 2:**
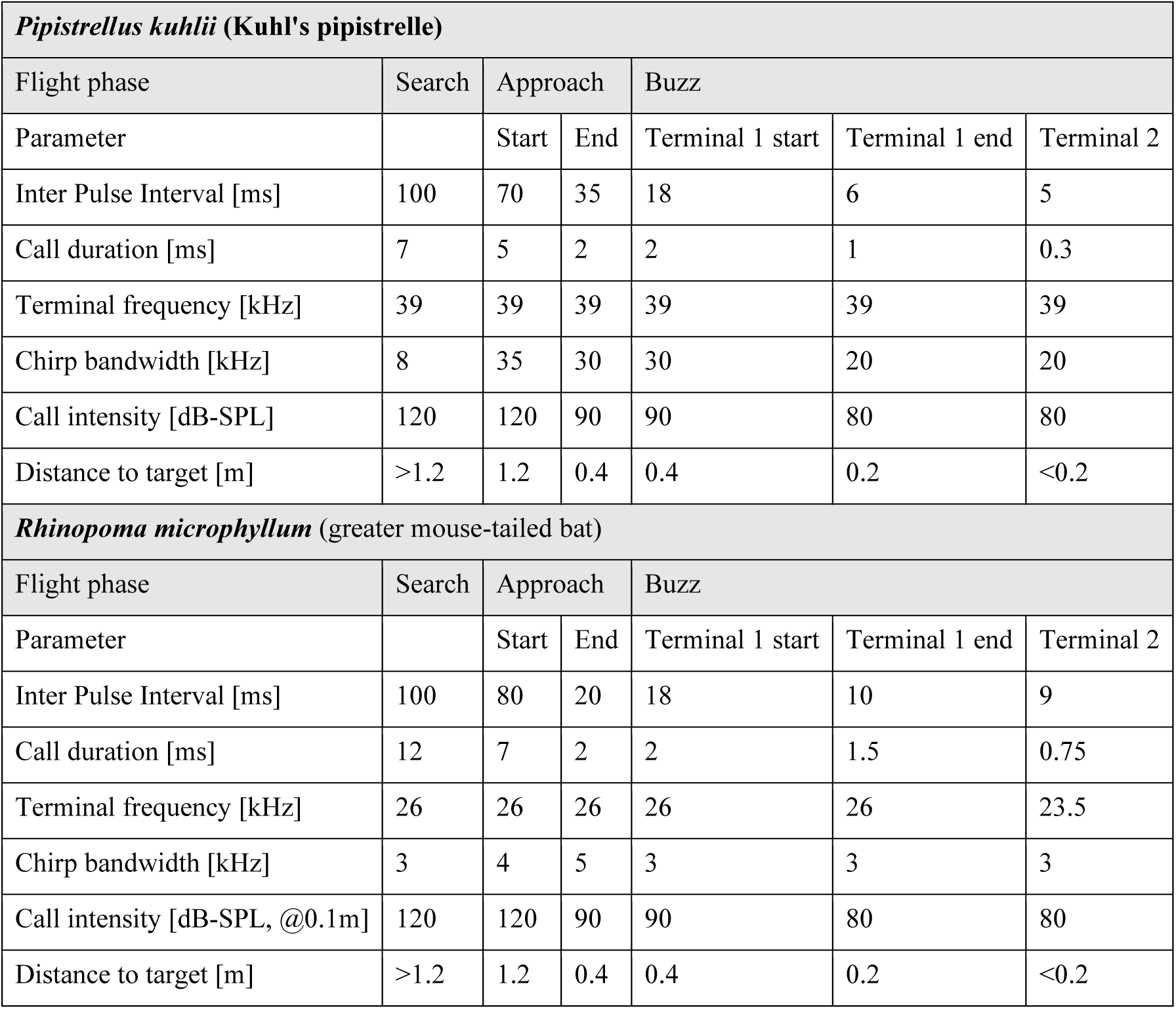
Echolocation parameters. The table presents the echolocation parameters of the two bat species we simulated during the specified flight phases (i.e., search, approach, buzz, and final buzz). In each phase, except for the search phase, in which the parameters remain constant, the parameters for each call are determined by the distance to the closest detected object.

While echolocation phases—search, approach, and buzz—are traditionally associated with prey capture, similar patterns have been documented in non-foraging tasks such as landing, obstacle avoidance, clutter navigation, and drinking^54–64^. In these contexts, bats modulate call duration and inter-pulse intervals according to object proximity, generating phase-like transitions even without prey. This supports the interpretation of phase structure as a general proximity-sensing strategy rather than a foraging-specific behavior. In our simulations, bats operated predominantly in the approach phase due to the cluttered cave environment—consistent with natural emergence behavior, where navigation dominates over open-space search. Accordingly, our use of echolocation phases in the model is biologically plausible across a range of sensory-guided tasks.

The bat densities we simulated, ranging from 1 to 100 bats per 3m^2^, reflect a wide range reported in field studies. Although bat colonies can be much larger than 100 bats, the maximal simulated density in our model (100 bats per 3 m²) resulted in bats flying in very close proximity, with an average nearest-neighbor distance of 0.27 meters. This density is higher than some of the most-dense reported bat aggregations, including studies on *Miniopterus fuliginosus*^49^, *Myotis grisescens*^65^, and *Tadarida brasiliensis*^4,50,66^, where bats emerge from the roost at rates of 15 to 500 bats per second, but fly with an average distance of 0.5 meters between individual bats.

We compared the performance of two FM echolocating insectivorous bat species: *Pipistrellus kuhlii (*PK) and *Rhinopoma microphyllum (*RM). PK bats emit wideband echolocation signals that are less prone to jamming than RM bats’ narrowband signal^15,67^, as wideband signals distribute energy across a broader frequency range and are thus more robust against interference^9,68^. Our findings show that PK signals slightly reduce jamming probability (by 9%) and improve wall detection. However, no significant differences in exit probabilities were noted between the two species.

Using a simulation allowed us to separate the effects of **acoustic interference (masking)** and **spatial interference (collision avoidance)** and revealed new insights into the sensorimotor strategy that could plausibly be used by real bats. The spatial interference reduced the probability of exiting the roost from 100% to 87%, while the acoustic masking further decreased it to 63%. Increasing call intensity had little effect on exit performance, although slightly improving it at high bat densities. When all bats increased their calling intensity, both desired echoes and masking signals intensified equally, resulting in only a marginal effect. This was tested by varying call intensity levels (100-130 dB SPL) in our simulations (Table 1), demonstrating that beyond a certain level (∼110 dB SPL), there is no further benefit in improving obstacle detection. These results align with previous studies that have drawn similar conclusions^7,24^.

Bats constantly adjust their flight speed to their surroundings^69–72^ and specifically when conspecifics are nearby^73^. Our study suggests that the optimal velocity for flying through a crowded roost ranges from 6 m/sec to 8 m/sec for densities of 2-100 bats/3m^2^. Exceeding this velocity-range led to a significant drop in exit probability due to a significant increase in wall collisions. We found that this speed did not depend on bat density in accordance with the observations of Theriault et al.^50^. Notably, the reported velocities of RM when exiting a cave^25^ and PK emergence velocity near the cave^74^ are close to the speed that appears optimal, based on our simulations.

We also tested the effects of wall and conspecific target strengths on navigation. Stronger wall echoes substantially improved exit probability and reduced obstacle collisions, despite slightly increasing masking of conspecific echoes (Supplementary Figure 3). In contrast, changes in conspecific reflectivity had minimal impact, likely because both desired and masking signals scaled similarly (Supplementary Figure 4). This result may also stem from our model’s assumption that bats slow down, but continue flying at the same direction following a collision with a conspecific.

Our basic model assumed that bats can distinguish between wall echoes and conspecific echoes, as demonstrated in previous studies ^75–77^. We suggest that this is a feasible assumption because echoes from cave walls are longer and exhibit distinct spectro-temporal patterns, whereas echoes from smaller objects, such as conspecifics, are shorter^47,78,79^. However, wall echoes reflected from conspecific calls might resemble those from the bat’s own calls in their amplitude and time-frequency characteristics ^20,73,80^. This led us to question how the misidentification of such echoes as obstacles might affect navigation. When unable to distinguish between these echoes, the simulated bats responded to all as if they were their own and thus mis-localized conspecific wall echoes. The confusion led to a substantial drop in exit performance, with only 15% of the bats successfully exiting compared to 82% under no-confusion conditions, at a density of 40 bats/3m^2^. At the same time, the collision rate increased markedly from 0.2 to 0.85 collisions per second. These results demonstrate the vital importance of echo discrimination for successful navigation, highlighting both the necessity of distinguishing between self and conspecific echoes and the classic challenge of detecting desired signals in noisy environments. There is a substantial evidence in the literature supporting the assumption that bats can recognize their own echoes and reliably distinguish them from those of conspecifics^68,75–77,81^.

Previous studies have also demonstrated that bats can aggregate acoustic information received sequentially over several echolocation calls, effectively constructing an auditory scene in complex environments^5,82–86^. Bats are also known to emit call sequences in groups, particularly when spatiotemporal localization demands are high. Studies have recorded sequences of 2-15 grouped calls, supporting the idea that grouping facilitates echo aggregation^83,87^. Accordingly, we tested how multi-call clustering process—which included grouping nearby reflectors, removing outliers, and estimating wall orientation based on these clusters—could assist bats in pathfinding, even under the assumption of full confusion. At bat densities of 1 to 40 bats/3m^2^ with masking, the multi-call clustering completely restored the collision rate with walls from 0.85 back to 0.2 collisions per second, and significantly improved the exit probability, raising it to 58%, although it did not entirely eliminate the impact of confusion. Our assumption of total confusion between echoes from a bat’s own calls and those from conspecifics, as well as our relatively simple clustering model, likely underestimates the true capabilities of real bats when flying in complex environments.

Navigation in bats involves processing complex sensory inputs and applying effective decision-making, often requiring an ability to switch strategies^88–94^. Bats possess a highly accurate spatial memory^82,90,94–96^, which is essential for tasks like long-distance migration^51^, homing^97^, and maneuvering in cluttered environments^95^. Additionally, they utilize acoustic landmarks to orient in total darkness^52^, occasionally rely on vision^91,92^, particularly at the cave edge where light is available, can passively detect echolocating peers, and perhaps eavesdrop on conspecifics’ echoes^23^. In this study we focused on whether echolocation alone is sufficient for one of the most difficult orientation tasks that bats perform – exiting a roost at high densities without prior knowledge of the roost’s shape, aside from the initial flight direction. Thus, our echolocation-only model, which was based on a five-call integration window during most simulations, probably underestimates real bats’ actual performance which also benefits from additional sensory input and can employ addition navigation strategies by sharing information between each other to coordinate and optimize the routes, such as manifested by swarming intelligence^33,98,99^.

Our model highlights the importance of considering sensory interference in animal behavior research and illuminates the impressive capabilities of echolocating bats. Additionally, the model showcases the value of simulations and establishes a framework for future studies on collective movement and swarming animals, and on robotics in complex environments.

## Methods

The simulated bats rely solely on echolocation to detect and locate obstacles and other bats by analyzing the sound waves they receive. They emit directional echolocation calls and receive the echoes reflected by roost walls and conspecifics, as well as the calls of conspecifics and the echoes returning from their calls. The bats adjust their flight trajectory and echolocation behavior based on the estimated location of the detected objects (range and angle), which deteriorates upon acoustic interference. The detection of the received signals is based on the mammalian gammatone filter bank receiver, under the assumption that bats can differentiate between the desired detected The sound intensity of the echoes generated by the bat’s own calls and those of its conspecifics are calculated using the sonar equation^7,108^ (pp. 196-198), as shown in Equation 1, geometrical relations are according to Supplementary Figure 5. The received levels of the masking calls are determined by using the Friis transmission equation^109^, as shown in Equation 2. All signal levels were simulated and reported in dB-SPL, referenced to 0.1 meters from the emitting bat. Bats are modeled acoustically as spherical reflectors with a fixed target strength of -23dB assuming reference distance 1 meter, reflecting sound isotropically. This approximates a sphere with a radius of 0.15 m, corresponding to the approximate wingspan of *Rhinopoma microphyllum* (RM) ^25,110^. While target strength can vary with wing posture and body geometry, we chose a representative value within the reported biological range for simplicity and model consistency. Our own measurement of a 3D-printed RM bat yielded a target strength of –32 dB, and a sensitivity analysis (Supplementary Figure 4) showed that model performance was only mildly affected across a wide range of target strengths (see Supplementary Figure 4). This supports the robustness of our approach to different sized bats. Walls are modeled as composites of individual reflectors placed 20 cm apart; each treated as a sphere with a 20 cm radius and a target strength of -22.5dB. For simplicity, in our model, the head is aligned with the body, therefore the direction of the echolocation beam is the same as the direction of the flight. The directivity of the calls and the received echoes is defined by the piston model^7,102^ with radii of 3 mm for the mouth-gap and 7 mm for the ear. The directivity is not directly influenced by velocity but follows behavioral dependent frequency changes. As the bat transitions from search to approach to buzz phases, it emits higher-frequency calls, leading to increased directivity. This shift coincides with a natural reduction in speed during the approach phase. Echo delays are calculated as the two-way travel time of the signals from the emitter to the target.

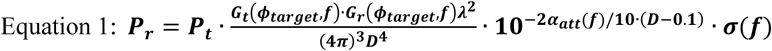

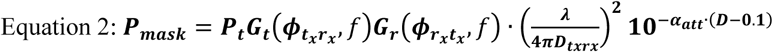

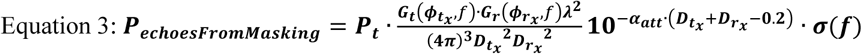

Where,

*P*_*r*_: level of the received signal [SPL]
*P*_*t*_ : level of the transmitted call [SPL]
*P*_*mask*_ : level of the masking signal as received by the bat [SPL]
*P*_*echoesFromMasking*_ : level of the echoes reflected by conspecifics and received by the bat [SPL]
G_t_(*ϕ*, *f*): gain of the transmitter (mouth of the bat, piston model), as a function of azimuth and frequency (*f*) [numeric]
*G*_*r*_(*ϕ*, *f*): gain of the receiver (ears of the bat, piston model) [numeric]
*ϕ*_*target*_ : the angle between the bat and the reflected object [rad]
D: distance between the bat and the target [m]
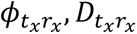 : the angle [rad], and the distance [m] between the transmitting conspecific and the receiving focal bat (from the transmitter’s perspective)
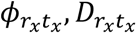 : the angle [rad], and the distance [m] between the receiving bat and the transmitting bat (from the receiver’s perspective)
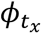 : the angle [rad], between the masking bat and target (from the transmitter’s perspective)
*α*_*att*_(*f*): atmospheric absorption coefficient for sound [dB/m]
*σ*(*f*): SONAR cross-section of the target [m^2^]
λ: The wavelength of the signal [m]

To maintain model simplicity, we did not incorporate Doppler effects in the echolocation model. While Doppler shifts can affect frequency perception, their impact on jamming and navigation performance is minimal in this context^111^. Moreover, the inter-individual random signals frequencies were larger than the expected Dopplers. In addition, the model does not assign echoes to earlier calls if their delays exceed the bat’s own Inter-Pulse Interval (IPI), and thus does not simulate pulse-echo ambiguity.

To model the detection process in the bat’s cochlea, we employed a monoaural filter bank receiver^47,112,113^ consisting of 80 channels, each with three components: (i) a gammatone filter of order 8, acting as a bandpass filter with center frequencies logarithmically scaled between 10kHz and 80kHz^7^; (ii) a half-wave rectifier; and (iii) a lowpass filter (Butterworth, fc=8kHz, order=6). Object detection and distance estimation are conducted using Saillant’s method^7,47,114^, based on the sum of detections in the active channels, see Figure 1C, D. Initially, a de-chirping process calculates the reference frequency-delay by detecting the peak in the response of each channel to the emitted call in a noise-free environment. Subsequently, the received signal, containing both desired echoes and masking sounds, passes through the filter bank. In each channel, all peaks above a threshold level are detected and time-shifted by the de-chirp reference. The detection threshold in each channel was set to the higher of two values: either 7 dB above the noise floor (0 dB-SPL) or the maximum received signal level minus 70 dB, thereby enforcing a dynamic range of 70dB. Peaks from all channels are aggregated in 5 µs windows and convolved with a Gaussian kernel with σ=5 µs. Output peaks that exceed the threshold level, set at 10% of the number of active channels, and fall within a time window of 100µs around the expected delay, are considered successful detections.

To evaluate the impact of acoustic interference, we conducted the detection procedure twice. The first, termed “interference-free detection”, comprised only the desired echoes, with white Gaussian noise at a level of 0 dB-SPL and without masking signals. The second, termed “full detection” comprised the desired echoes, Gaussian noise, and the masking signals. Detected echoes in the full detection were defined by the strongest peak within a four-millisecond window (three milliseconds before and one millisecond after, accounting for forward and backward masking ^24,115–117)^ detected above the threshold within 100µs of the interference-free detections. If the detected peak in the full detection condition was delayed by more than 100 µs compared to the interference-free case, it was defined as a miss-detection. Peaks with smaller timing shifts were considered **detection with timing errors. Jammed echoes** were defined as echoes that were detected under the interference-free condition but not detected under the full detection condition. The **jamming probability** was calculated as the ratio of jammed echoes in the full detection condition to the detected echoes in the interference-free condition.

After detection, the bat estimates the range and the Direction of Arrival (DOA) of the reflecting objects. The range is determined by the delay of the detected echo, including any errors derived from the filter-bank process in the “full detection” process (i.e., including all masking signals).^7,110,113^. The direction is not explicitly estimated through binaural processing. Instead, based on previous studies ^115,118^, we assumed that bats can estimate the direction of arrival with an angular error that depends on the Signal-to-Noise Ratio (SNR) and the angle. The inputs to this process include the peak level of the desired echo, the noise level, and the level of acoustic interference. The output is the estimated direction of arrival with a random error applied based on the SNR. At an angle of 0° and an SNR of 10 dB, the standard deviation of the error is 1.5° ^119^ and ^7^ (Equation 4), with the error capped at a maximum of 3° in our model.

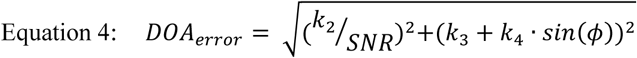

where, k₂, k₃, and k₄ are constants chosen to produce a DOA error consistent with the range described above.

To evaluate the impact of the assumption that bats can distinguish between echoes caused by their own calls and those caused by other bats (i.e., conspecifics’ reflectors), we tested an alternative model in which the simulated bats treat all echoes reflected from walls as if they have originated from their own calls The distance to reflectors of conspecifics’ calls is estimated based on the time difference between the echo and the bat’s last call. The direction of arrival is estimated by the angle between the bat and the physical reflector, with an added random error (the same process used for their own echoes).

In real bats, spatial processing in the brain involves integrating auditory and spatial information over time to construct a coherent map of the environment ^5,68^. This neural computation is crucial for navigation and prey detection in complex environments. To examine whether spatial integration mitigates the confusion problem, we added a ‘multi-call clustering’ module that was based on the sensory information obtained within a one-second memory window. The clustering comprised the following steps: (i) clustering all detections in memory into groups with a maximum internal distance of 10 cm; (ii) reconstructing the estimated walls positions and directions based on the average of clusters that include at least two detections (rather than relying on single reflections); and (iii) identifying openings between reconstructed wall edges ranging from 0.5 to 2.25 meters in width, see Supplementary Figure 1 and Supplementary Figure 2B. The model assumes that bats store echo locations in an allocentric x-y coordinate system, transforming detections from a local to a global spatial framework. Collision avoidance is based not only on the integrated spatial representation but also on immediate echoes from the last call (prior to clustering), including potential uncorrected false detections and localization errors, which are independently processed for real-time evasive maneuvers.

### Statistical analysis

Statistical analysis and the roost-exit model were conducted using MATLAB© 2023a.

Tests were performed with a significance level of 0.05. For each simulated scenario, we examined the effect of the various parameters on exit probability, time-to-exit, collision rate, and the jamming probability, using Generalized Linear Models (GLMs). The GLM tests were executed with MATLAB built-in function **‘fitglm()’**. Probability variables (such as exit and jamming probabilities) were treated as binomially distributed; rate variables (such as collision rate) were treated as Poisson distributed, and all other variables were considered normally distributed. Unless otherwise stated, all explaining variables were set as fixed factors. All statistical analyses, including the statistical test and the corresponding sample sizes, are described throughout the text and summarized in Table 1. Standard errors are calculated across all individuals within each scenario, without distinguishing between different simulation trials.

## Supporting information

Supplementary file- a video example of 10 bats

Cover letter

Authors Response

## Author Contributions

**O.M** - Software, Formal analysis, Data acquisition, Validation, Visualization, Methodology, Writing - original draft, Writing - review and editing. **Y.Y** - Conceptualization, Resources, Supervision, Funding acquisition, Validation, Investigation, Methodology, Project administration, Writing - review and editing

## Data availability

All data and codes generated during this study are included in the manuscript and supporting files. Source code files have been uploaded with a Graphical User Interface and a readme file for explanation. Data are available at zenodo and github:

https://zenodo.org/records/16992617 (link)
https://github.com/omermazar/Colony-Exit-Bat-Simulation/tree/main (link)

## Supplementary

Supplementary Movie 1

[link](https://drive.google.com/drive/folders/1sp-Zh9NfQqFLD6eB83Jg3WmqHEeW88Y7)

**Supplementary Figure 1:**
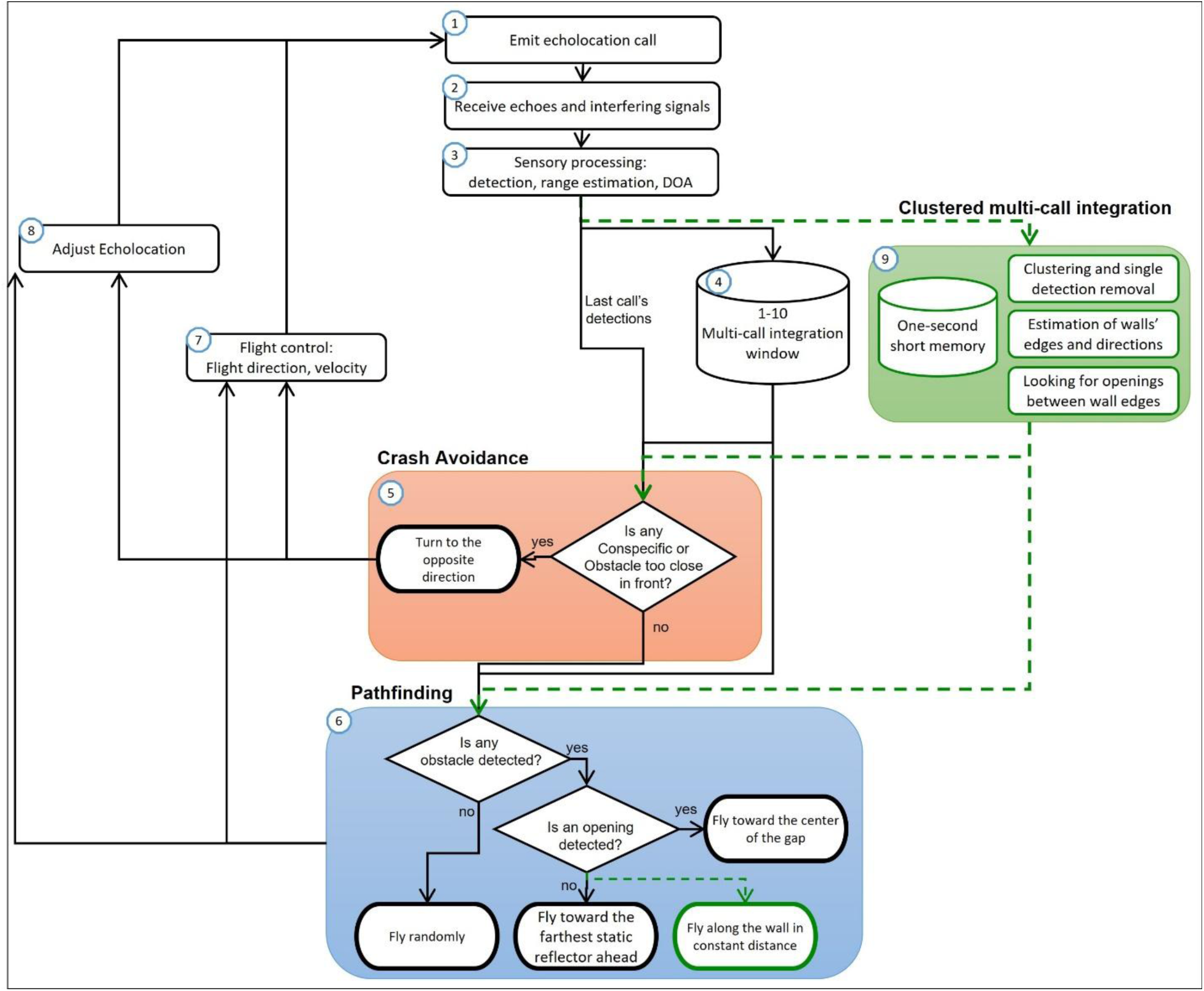
Decision-making in echolocation-based pathfinding.

This diagram illustrates the sensorimotor decision-making process based solely on echolocation. The process starts with the emission of an echolocation call (1) and the reception of echoes and interfering signals (2), followed by sensory processing for detection, range estimation, and direction of arrival (DOA) (3). After integrating detections over a 1–10 call window (4), the bat engages in **crash avoidance** (5) by evaluating the proximity of conspecifics and obstacles directly ahead. If either is too close, the bat turns in the opposite direction of the detected obstacle, by applying maximum angular velocity away from it (e.g., if the obstacle is on the right, the bat turns left). If no immediate threat is detected, the bat proceeds to **pathfinding** (6). During pathfinding, it checks for obstacles and, if an opening is detected, flies toward the gap’s center. Without the optional **multi-call clustering process** (green), the bat simply integrates detections and flies toward the farthest detected obstacle, interpreting it as a wall edge. If the multi-call clustering is included (9), a one-second short memory aids in clustering detections, estimating wall edges, and identifying openings, while also allowing the bat to follow walls at a constant distance. Throughout, the bat continuously adjusts echolocation parameters (8) and controls flight direction and velocity (7) based on ongoing sensory information and decision-making.

**Supplementary Figure 2A:**
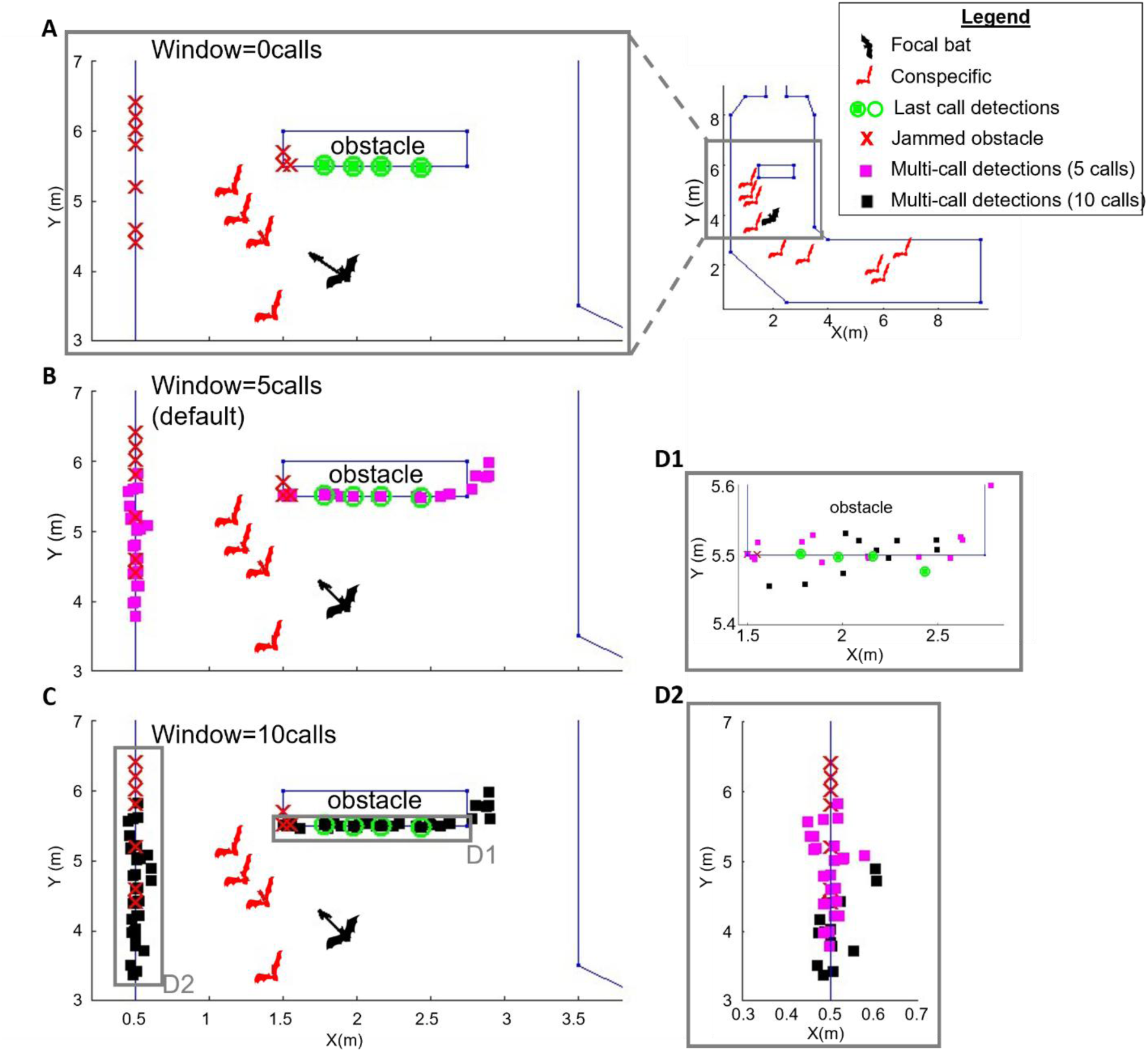
Multi-Call Integration.

This figure demonstrates the effect of multi-call integration under non-confusing conditions. The upper-right panel shows the position of the focal bat (black) and nine conspecifics (red) within the roost corridor, with a zoomed-in view of the gray rectangle provided in Panels A–C. **(A)** When the integration window is set to zero calls (no memory), the bat relies solely on the latest call. Green circles and squares represent detected reflectors, while red Xs indicate missed (jammed) detections. Notably, the left wall of the corridor remains undetected due to jamming. **(B, C)** Increasing the integration window to five calls (magenta squares) and ten calls (black squares) accumulates detections from prior calls, improving coverage of the environment. In this basic integration model, each detection is treated independently, without clustering. **(D1, D2)** Magnified views of the grey regions indicated in Panel C, comparing detections across 0, 5, and 10-call windows (green, magenta, and black, respectively), illustrating how extended memory improves detection robustness. Note that the X-Y aspect ratios in D1 and D2 differ from the main panels to enhance visibility of spatial distributions.

**Supplementary Figure 2B:**
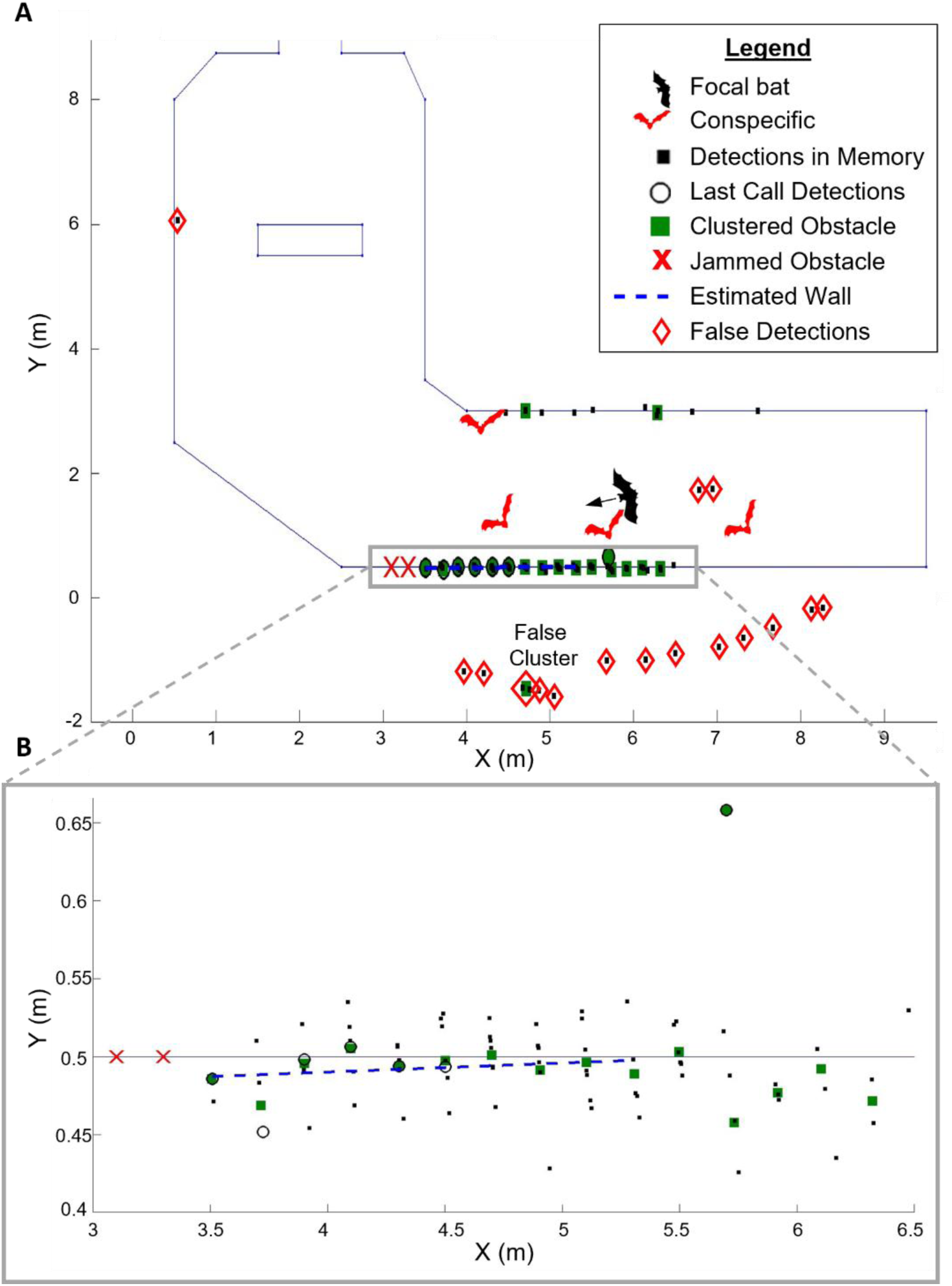
Multi-Call Clustering Example.

This figure illustrates the multi-call clustering algorithm under full-confusion conditions. (A) A focal bat (black) and four conspecifics (red) are shown in the lower corridor. (B) A zoom-in of the gray rectangle in (A). Black ovals represent detections from the last call; red X’s indicate jammed echoes; black squares represent all detections stored across the integration window (before clustering), each subject to localization error. When not applying multi-call clustering – the bat would rely on all of these dots as reflectors. Under full confusion, the bat cannot distinguish self-echoes from conspecific echoes, leading to false detections (red diamonds). Detections are clustered when a reflector is detected twice or more within a 10 cm radius (green squares). The clustered reflectors are used to estimate wall directions (blue dashed line) and detect possible gaps (not shown). As a result of to the multi-call clustering algorithm, most false detections are removed as outliers, except for one erroneous cluster (Panel A). Collision avoidance maneuvers are based on both the clustered obstacles and the raw detections from the latest call (empty black ovals).

**Supplementary Figure 3:**
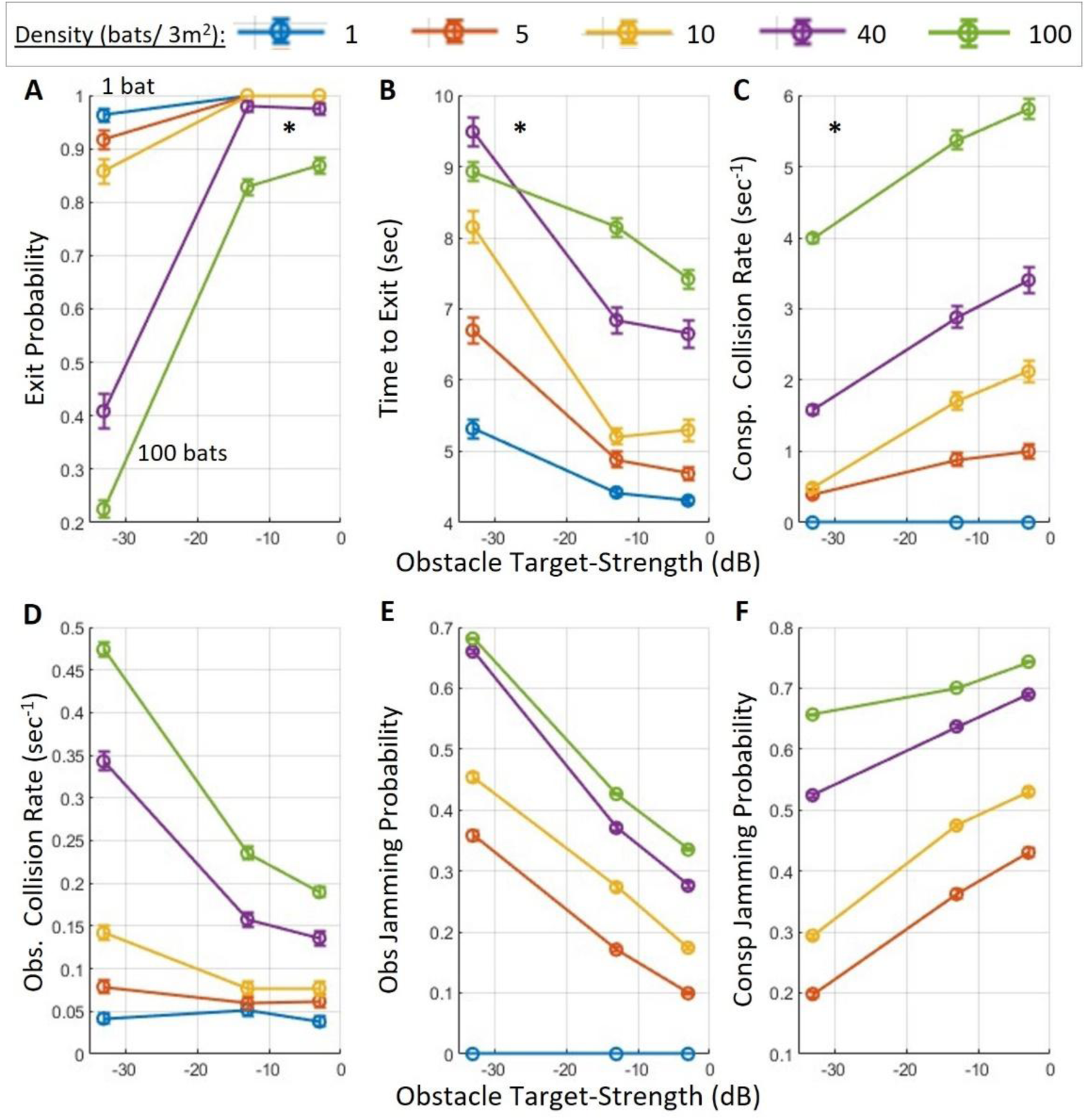
Sensitivity of exit performance to obstacle target strength.

This figure shows how changes in the acoustic target strength of the cave walls affect navigation performance across five bat densities (1, 5, 10, 40, and 100 bats/3 m²). Target strength values ranged from –33 dB to –3 dB, corresponding to spherical reflectors with approximate radii from 0.05 m to 1.5 m. Overall, increasing obstacle target strength significantly influenced exit performance, primarily by reducing the probability of obstacle jamming and thereby improving detection. **(A) Exit Probability** increased with obstacle target strength across all densities, with a maximal increase of 64% for a density of 100 bats (p << 10⁻¹⁰, t = 28.5, DF = 8157, GLM)**. (B) Time to Exit** decreased significantlywith increasing obstacle target strength, with a maximal reduction of approximately 2.8 seconds at a density of 10 bats (p << 10⁻¹⁰, t = –22.2, DF = 6920, GLM). **(C) Conspecific Collision Rate** increased slightly with stronger obstacle echoes (p << 10⁻¹⁰, t = 27.6, DF = 8157, GLM). **(D) Obstacle Collision Rate** decreased significantly with increasing target strength (p << 10⁻¹⁰, t = –10.7, DF = 8157, GLM), reflecting better detection of walls and structures. **(E) Obstacle Jamming Probability** decreased consistently (p << 10⁻¹⁰, t = –19.8, DF = 8157, GLM). **(F) Conspecific Jamming Probability** increased with obstacle target strength (p << 10⁻¹⁰, t = 27.6, DF = 8157, GLM).

These results suggest that stronger wall echoes improve environmental awareness at the cost of slightly increased masking of conspecific echoes. Despite this, the overall performance—particularly exit probability and reduced obstacle collisions—improves significantly.

In all panels, circles represent means and bars represent standard errors. The error bars are present but very small due to the large number of simulation repetitions, and thus may not be visually noticeable at the plotted scale. See Table 1 for the number of simulated bats.

**Supplementary Figure 4:**
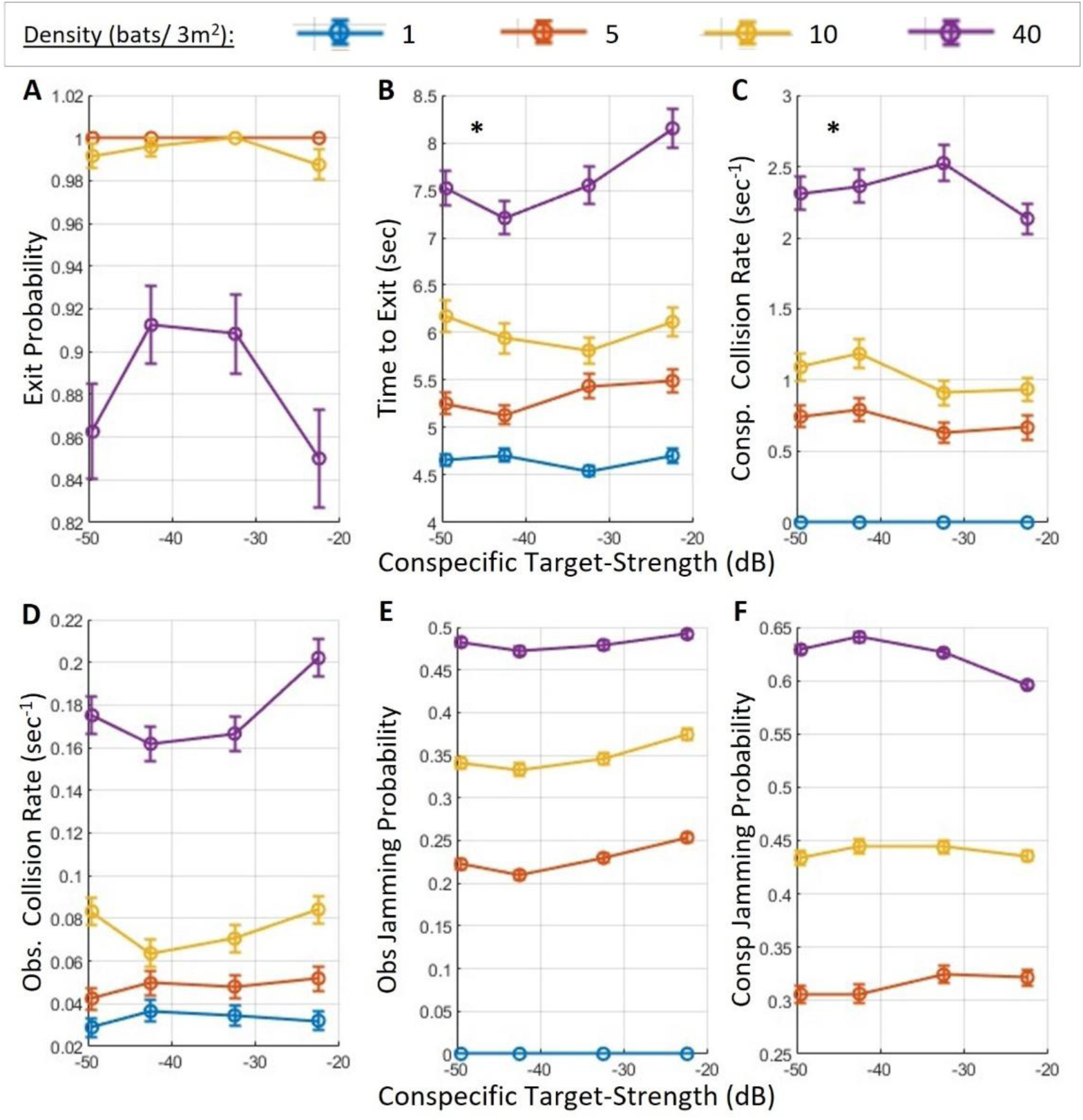
Sensitivity of exit performance to conspecific’s target strength.

This figure shows how changes in the acoustic target strength of conspecifics affect navigation performance across four bat densities (1, 5, 10, and 40 bats/3 m²). Overall, our results indicate that target strength has a relatively minor impact on performance, likely because it affects both desired echo signals and masking signals equally. Interestingly, this analysis also suggests that our model is more sensitive to the bat’s response to nearby conspecifics than to the physical collision impact itself. **(A)** Exit probability was not significantly affected by conspecific target strength (p=0.28, t=-1.09, DF=5757, GLM, see details in Table 1). Note that the performance curves for densities of 1 and 5 bats overlap almost completely. **(B)** Time-to-exit increased with target strength at high density, with a maximal effect size of ∼1 second at 40 bats (p = 0.003, t = 3.02, DF = 5578). **(C, D)** Collision rates with conspecifics decreased significantly with stronger target strength (p = 0.0002, t = –3.7, DF = 5757), while collisions with obstacles remained statistically unchanged (p = 0.23, t = 1.18, DF = 5757). **(E, F)** Jamming probability was not significantly affected for either conspecific or obstacle echoes (p = 0.6, t = –0.51, DF = 4762; p = 0.19, t = 1.31, DF = 5757, respectively). This aligns with the notion that both useful and interfering signals scale similarly with target strength. Importantly, the probability of detecting a conspecific located within 1 meter increased substantially with higher target strength, improving from 25% to 43% at 40 bats (p < 10⁻¹⁰, t = 6.45, DF = 4162).

In all panels, circles represent means and bars represent standard errors. The error bars are present but very small due to the large number of simulation repetitions, and thus may not be visually noticeable at the plotted scale. See Table 1 for the number of simulated bats.

**Supplementary Figure 5:**
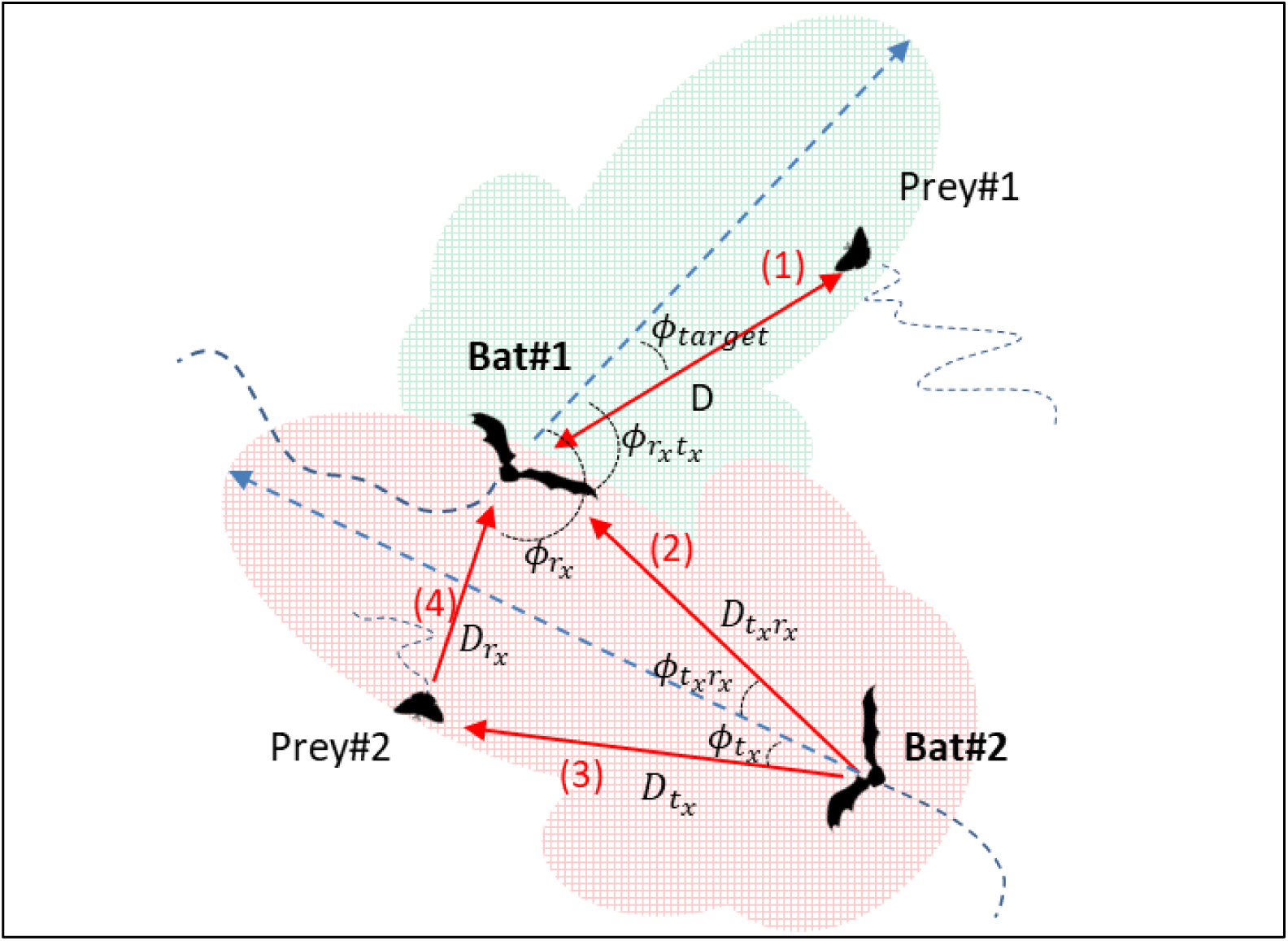
Angles and distances for two bats and two reflecting objects.

Bat1 receives a reflected echo from Prey1 or a stationary obstacle located at a distance of D from it, with an angle ϕ_target_ relative to its flight direction (red arrow 1). Prey1 is also within the detection range of Bat1, depicted by the green shaded piston area. Bat1 also receives masking sounds from Bat2. The echolocation signals emitted by Bat2 arrive at the ear of Bat1 at an angle ϕ_txrx_ relative to its flight direction and from a distance of D_txrx_ (red arrow 2). Additionally, the echolocation signals of Bat2 are reflected by Prey2, before being received by Bat 1. These reflected signals act as masking signals at a relative angle of angle ϕ_rx_, and from a distance of D_rx_ from Bat1.

## Notes

https://github.com/omermazar/Colony-Exit-Bat-Simulation

### Competing Interest Statement

The authors have declared no competing interest.

### Summary of Updates

We have carefully addressed all remaining issues and clarified the points raised in the latest round of reviews. Below, we provide detailed responses to each comment, noting the revisions made in the manuscript.

### Funding

- Israel Science Foundation: 331/22

