## Supplementary material for "How bats exit a crowded colony when relying on echolocation only - a modeling approach": Cover letter

Dear Editor,

We are pleased to **resubmit** our manuscript entitled "***How bats exit a crowded colony when relying on echolocation only - a modeling approach***" for consideration by ***eLife***.

Bats flying in dense groups in confined spaces face some of the most difficult sensory tasks with a need to detect weak signals (echoes) amidst a cacophony of noise (conspecific calls and echoes). In this study, we build upon our previously published research in *eLife* (Mazar and Yovel 2020), where we introduced an initial sensorimotor model of bat echolocation. Since then, we have significantly improved this model by adding temporal integration and navigational components. This allowed us to simulate more challenging scenarios and explore additional highly challenging and applicable scenarios as we do in this study. This advancement is particularly relevant as many sensory systems, both biological and artificial, face similar challenges of detecting and discriminating weak signals in noisy environments.

Our updated model demonstrates that the bat echolocation system is remarkably resilient to jamming interference, even when flying in extremely dense groups and in highly cluttered environments, such as narrow caves. By mimicking the echolocation behavior of two bat species under such high-density conditions, our model highlights how strategies routinely applied by echolocating bats, overcome acoustic jamming and enable effective pathfinding even in complete darkness. The modeling approach reveals many advantages as it allows us to fully simulate the sensory input from the bat's point of view. To ensure biological plausibility, we intentionally used a simple model that underestimates the bats' actual abilities.

In this revised version, we have carefully addressed all comments raised by the reviewers. This includes providing clearer explanations of model assumptions and parameters (e.g., multi-call integration and clustering, echo discrimination, and behavioral parameters), clarifying figures and their captions, and adding Supplementary Figures for more detailed explanations. We also revised several parts of the Abstract and Discussion to better emphasize key results—particularly the strong impact of multi-call integration on navigation performance. Minor textual improvements were made throughout to enhance clarity without altering the substance of the findings. These changes aim to improve transparency and reproducibility, in line with *eLife*'s standards.

This research addresses a fundamental challenge in active sensing systems — how organisms detect and process signals under multi-source interference — offering insights that advance understanding in biology, bio-inspired design, and autonomous robotics, particularly in noisy, dynamic settings

This manuscript is original, unpublished, and not under consideration elsewhere. We have no conflicts of interest to disclose.

We thank the reviewers and editors for their constructive feedback, which helped us improve the manuscript.

Thank you for your consideration of this manuscript.

Ethical issues: The study did not involve live animals

Funding: This project was funded by Israel Science Foundation 331/22 (Y.Y)

Sincerely,

Omer Mazar and Yossi Yovel
